# The AAA+ ATPase RavA-ViaA complex sensitizes *Escherichia coli* to aminoglycosides under anaerobic low energy conservation conditions

**DOI:** 10.1101/2022.01.13.476284

**Authors:** Jessica Y. El Khoury, Jordi Zamarreño Beas, Allison Huguenot, Béatrice Py, Frédéric Barras

## Abstract

Aminoglycosides have been used against Gram-negative bacteria for decades. Yet, uncertainties remain about various aspects of their uptake mechanism. Moreover their killing efficiency is well known to vary as a function of growth conditions and types of metabolism used by the targeted bacterium. Here we show that RavA, an AAA+ ATPase from the MoxR subfamily, associated with its VWA-containing partner, ViaA sensitize *E. coli* to lethal concentrations of AG, including gentamycin (Gm) and tobramycin, but not of antibiotics of other classes. We show this sensitizing effect to be due to enhanced Gm uptake in a proton motive force dependent manner. We evaluated the influence of RavA ViaA throughout a series of growth conditions, including aerobiosis and anaerobiosis. This led us to observe that the sensitizing effect of RavA ViaA varies with the respiratory chain used, i.e. RavA ViaA influence was prominent in the absence of exogenous electron acceptor or with fumarate, i.e. in poor energy conservation conditions, and dispensable in the presence of nitrate or oxygen, i.e. in high level of energy conservation. We propose RavA ViaA to be able to sense energetic state of the cell and to be used under low energy conditions for facilitating uptake of chemicals across the membrane, including Gm.

**Author Summary:** Antibiotic resistance is a major public health, social and economic problem. Aminoglycosides are known for their high efficiency against Gram-negative bacteria but their use is restricted to life threatening infections because of their nephrotoxicity and ototoxicity at therapeutic dose. Elucidation of AG sensitization mechanisms in bacteria will allow the use of a decreased effective dose of AGs. Here we identified new molecular actors, RavA and ViaA, which sensitize *E. coli* to AG under anaerobiosis. RavA belongs to the AAA+ ATPase family while ViaA bears a VWA motif. Moreover we show here that the influence of RavA ViaA on AG sensitivity varies with growth conditions and respiratory metabolism used by *E. coli*. This is a significant step forward as anaerobiosis is well known to reduce antibacterial activity of AG. This study emphasizes the crucial importance of the relationships between culture conditions, metabolism and antibiotic resistance.

## Introduction

Antibiotic resistance is an important biomedical problem that challenges the ability to treat bacterial infections [1]. Modulation of intracellular concentrations of antibiotics is one of the most frequent processes leading to resistance whether it is due to limiting antibiotic entry, or increasing efflux [2]. The aminoglycosides (AGs) class of antibiotics targets the ribosome, leading to mistranslation and eventually cell death. AGs comprise kanamycin, tobramycin, gentamycin (Gm), neomycin, amikacin and streptomycin and are commonly used worldwide, thanks to their high efficacy and low cost. The mechanism of action of AG has been studied for decades. Yet, uncertainties remain on key mechanistic issues, such as mode of killing and penetration into cells. For instance, although ribosome as the primary targets of AG has been established for years, several studies have in the last decade challenged the idea that ribosome targeting was the actual reason for AG toxicity, giving to reactive oxygen species an equally important contribution to bacterial death [3–6]. Another issue under debate pertains to the mechanism of entry of AG and various non-exclusive mechanisms have been described. The current model [7,8] is that uptake starts with the entry of a small amount of AG through proton motive force (*pmf*)-dependent mechanisms [9–11] or other transport systems [12–14] priming mistranslation by the ribosome, which leads to membrane damage by incorporation of mistranslated proteins and a second wave of massive AG uptake [9,15]. The *pmf* is especially important during the first phase of uptake [16], while the second phase occurs in response to impaired translation. *pmf* is produced by the activity of electron transfer chains arising within respiratory complexes and we previously showed that maturation of respiratory complexes is directly impacting AG uptake efficiency and level of resistance [4].

The AAA+ ATPases are widely used as part of macromolecular machines [17]. AAA+ ATPases share structural features such as forming oligomers or coupling ATPase hydrolysis with remodeling of substrates. AAA+ ATPases have been classified in different 7 clades [17]. Actual physiological role of several AAA+ ATPase remains to be elucidated, in particular within members of the MoxR subfamily, which is part of the AAA+ ATPase clade 7 [17]. Studies on different members of the MoxR family indicate that they could fulfill chaperone-like roles assisting assembly of protein complexes [18]. A hallmark of this type of AAA+ ATPase is that they interact with proteins containing the von Willebrand factor (VWA) domain, a domain that mediates protein-protein interactions. Among the MoxR family we are interested in the protein **r**egulatory **v**ariant **A** (RavA) with its VWA containing partner ViaA. RavA ViaA (RV) complex was reported to interact physically with the respiratory complex fumarate reductase (Frd) but the role for RV in fumarate respiration remains unclear [19]. Briefly, Frd allows *E. coli* to use fumarate as terminal electron acceptor when oxygen is lacking. Frd complex is made of four subunits, including the cytosolic soluble FrdA and FrdB subunits and the membrane spanning FrdC and D subunits. FrdB hosts a flavin adenine dinucleotide (FAD) and FrdA three iron-sulfur (Fe-S) cluster cofactors. FrdB is a menaquinone (MQ) oxidoreductase, which relays electron from MQH2 to FrdA active site that reduces fumarate into succinate [20]. RV was found to decrease Frd activity and it was proposed that RV complex might participate to the multi-step assembly process leading to functional Frd. A link between RV and anaerobic respiratory metabolism is also supported by the fact that *ravA-viaA* genes form an operon regulated by the anaerobic sensing transcriptional regulator Fnr [19]. Other potential partners/subtrates of RV are some members of the Nuo respiratory complex and of proteins of the iron-sulfur cluster biogenesis machinery Isc [21].

RV activity appears to be important for sustaining stress imposed by the presence of sublethal concentration of AGs, both in *Escherichia coli* and *Vibrio cholerae* [21–23]. Precisely, *E. coli* strains lacking *ravA*, *viaA* or both reached a higher final cell density than WT, in rich medium supplemented with sub-lethal concentrations of AGs [21]. Importantly, these observations were collected with *E. coli* strains growing under aerobic conditions, i.e. conditions in which neither Fnr transcriptional activating nor fumarate reductase activities are expected to intervene.

In the present work, we tested whether RV plays a role in *E. coli* sensitivity to lethal doses of AG. We carried out analyses both under anaerobic and aerobic growth conditions. Our study establishes RavA and ViaA as AG toxicity enhancers and found this activity to depend upon *pmf*. Unexpectedly, importance of RV in mediating AG toxicity varies with the growth conditions and nature of the electron acceptor provided to *E. coli* for respiring. We propose RV to connect the energetic status of the cell and AG uptake.

## Results

### RavA and ViaA sensitize *E. coli* to Gm under anaerobic fumarate respiratory conditions

The *ravA-viaA* operon is activated under anaerobiosis by the Fnr transcriptional activator [19]. Moreover, previous studies have pointed out a link between RV complex and Frd fumarate reductase activity [19]. Therefore we tested the effect of RavA ViaA on AG sensitivity of *E. coli* grown under anaerobic fumarate respiratory conditions. First, we investigated the importance of the fumarate respiration for Gm survival. We found that in a time-dependent killing experiment using a concentration of Gm equivalent to 2X MIC (16 μg/mL), the Δ*frdA* mutant exhibited increased resistance to Gm compared to the WT strain (Fig. 1A). Under anaerobic respiration, menaquinone is predicted to be a prominent electron carrier [24]. Therefore we tested the Δ*menA* mutant, which lacks an 1,4-dihydroxy-2-naphthoate octaprenyltransferase involved in menaquinone biosynthesis. Δ*menA* mutant exhibited same level of Gm resistance as Δ*frdA mutant* (Fig. 1B). Fumarate respiration might involve the NADH:quinone oxidoreductase (Nuo) or the anaerobic glycerol-3-phosphate dehydrogenase (GlpA, B & C) complexes as primary electron donors [25]. Accordingly, we tested the effect of mutations in both of these complexes, e.g. *nuoC* and *glpA*, on Gm killing. Neither Δ*nuoC* nor Δ*glpA* mutation, isolated or in combination, altered the level of Gm sensitivity (S1 Fig.). These results showed that functional menaquinone and fumarate reductase, as electron carrier and terminal reductase respectively, are required for Gm killing. Then we tested the Δ*ravA-viaA* mutant in a time-dependent killing experiment, using a concentration of Gm equivalent to 2X MIC (16 μg/mL). We found Δ*ravA-viaA* mutant exhibited increased resistance to Gm compared to the WT strain (Fig. 1A). Last, combining Δ*ravA-viaA* and Δ*frdA* mutations showed no additive effect (Fig. 1A). Altogether these results indicated that in fumarate respiratory conditions RavA ViaA sensitize *E. coli* to Gm in a FrdA-dependent mechanism.

**Fig 1.**
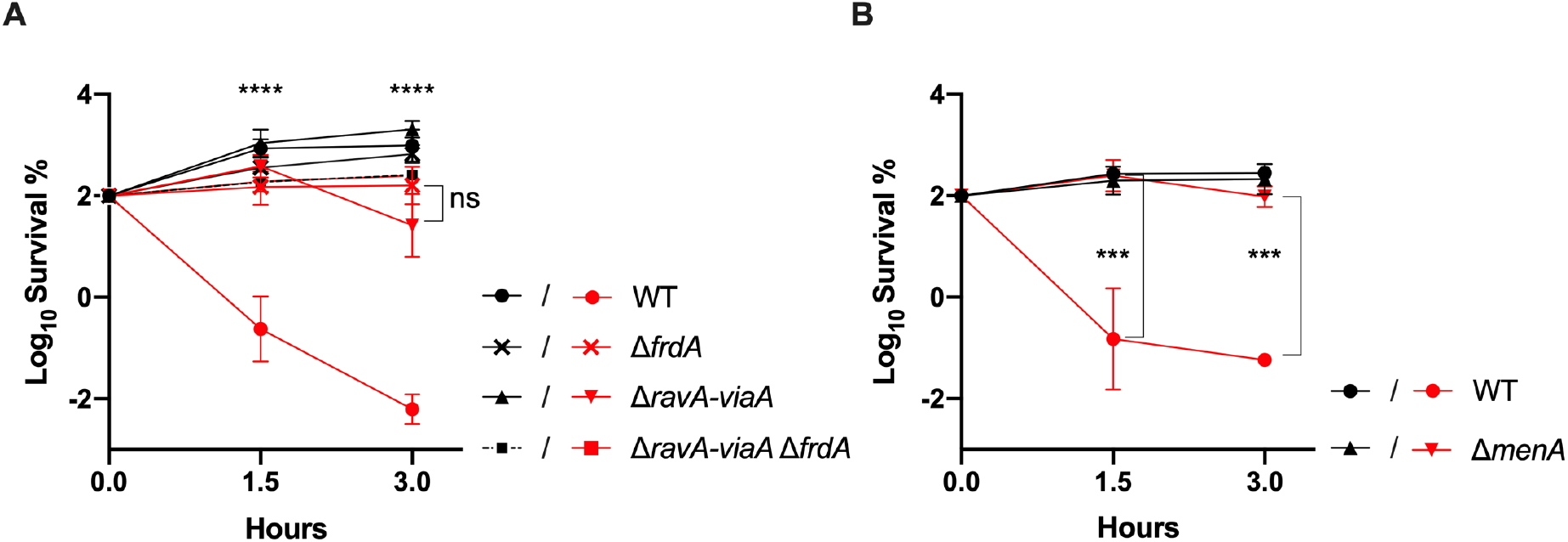
The *ravA-viaA* operon sensitizes fumarate respiring *E. coli* to gentamycin under fumarate respiration. (**A & B**) Survival of WT (FBE051), Δ*ravA-viaA* (FBE706), Δ*frdA* (FBE790), Δ*ravA-viaA* Δ*frdA* (FBE831) and Δ*menA* (FBE501) strains after Gm treatment. Cells were grown anaerobically in LB supplemented with fumarate at 10 mM and then Gm was added at 16 μg/mL. The survival values after 1.5 and 3 hours of treatment are represented. Black and red lines are for untreated and Gm-treated bacteria, respectively. For Δ*ravA-viaA* Δ*frdA* (FBE831) strain, lines of treated and untreated cells are overlapping. Survival measured by CFU per mL, was normalized relative to time zero at which Gm was added (early log phase cells; ~5×10^7^ CFU/mL) and plotted as Log_10_ of % survival. The Minimal Inhibitory Concentration (MIC) value of Gm was 8 μg/mL in LB medium supplemented with fumarate at 10 mM for the four strains, WT, Δ*ravA-viaA*, Δ*frdA and ΔmenA*, grown anaerobically. Values are expressed as means of at least 3 biological replicates and error depict standard deviation. One-way ANOVA tests followed by Sidak’s multiple comparaison tests were performed to compare at each time point (1.5 and 3 hours) the treated WT to each of the treated mutant, in (A) asterisks were similar for the three mutants and therefore were represented once for all (*** adjusted *p*Value = 0.0002 & **** adjusted *p*Value < 0.0001).

### RavA and ViaA are not needed to grow under fumarate respiration

Previous analysis showed that RV exerted, if anything, a slight negative effect on FrdA enzymatic activity [19]. We therefore tested if RV provided a growth advantage to strains growing using fumarate respiration. Both WT and Δ*ravA*-*viaA* strains were grown in LB medium supplemented with 10mM fumarate. Colony-forming unit (CFU) counting showed no difference between Δ*ravA*-*viaA* mutant and WT strain (S2.A Fig). Next, both strains were mixed at a 1:1 ratio, and grew together in M9-glycerol and fumarate under anaerobic conditions for 48h. Competitive index was determined by counting Δ*ravA*-*viaA* CFU, using their kanamycin resistance phenotype, and WT strain CFU at t_0_ and t_48_. The competitive index (CFU_mutant_/ CFU_wt_)t_48_/(CFU_mutant_/ CFU_wt_)t_0_ gave a median value of 1.3, revealing no growth advantage of one strain over the other (S2.B Fig). Altogether these results showed that *in vivo* RavA and ViaA do not influence growth under fumarate respiration and presumably do not modulate FrdA enzymatic activity to a large extent, if at all.

### RavA and ViaA do not sensitize *E. coli* grown under nitrate respiration to Gm

Nitrate can be used as a terminal acceptor yielding to the most energetically favourable anaerobic respiratory chain in *E. coli* [26]. Therefore, we tested if RavA and ViaA were able to sensitize *E. coli* to Gm under nitrate respiration. Nitrate reductase NarGHI is used under anaerobic conditions and is the major enzyme during nitrate respiration. First, we established the MIC value of Gm in LB medium supplemented with NaNO_3_ at 10 mM and 0.2 % glycerol and found it was 8 μg/mL for the WT, Δ*ravA-viaA*, Δ*narG* and Δ*ravA-viaA* Δ*narG* strains. Next, in these conditions we performed a time-dependent killing assay with Gm at 16 μg/mL. The Δ*ravA-viaA* mutant was as sensitive as the WT strain (Fig. 2), while Δ*narG* mutation made *E. coli* less susceptible to Gm comparing to the WT. These results indicate that the sensitization of *E. coli* to Gm under nitrate respiration is dependent upon NarG but independent of RavA-ViaA. Note that combining Δ*ravA-viaA* and Δ*narG* had a slight additive effect as the resulting strain showed an even higher level of resistance than the Δ*narG* single mutant (Fig. 2) suggesting that preventing respiration of the exogenously added electron acceptor, nitrate, permitted RV to sensitize *E. coli* to Gm.

**Fig 2.**
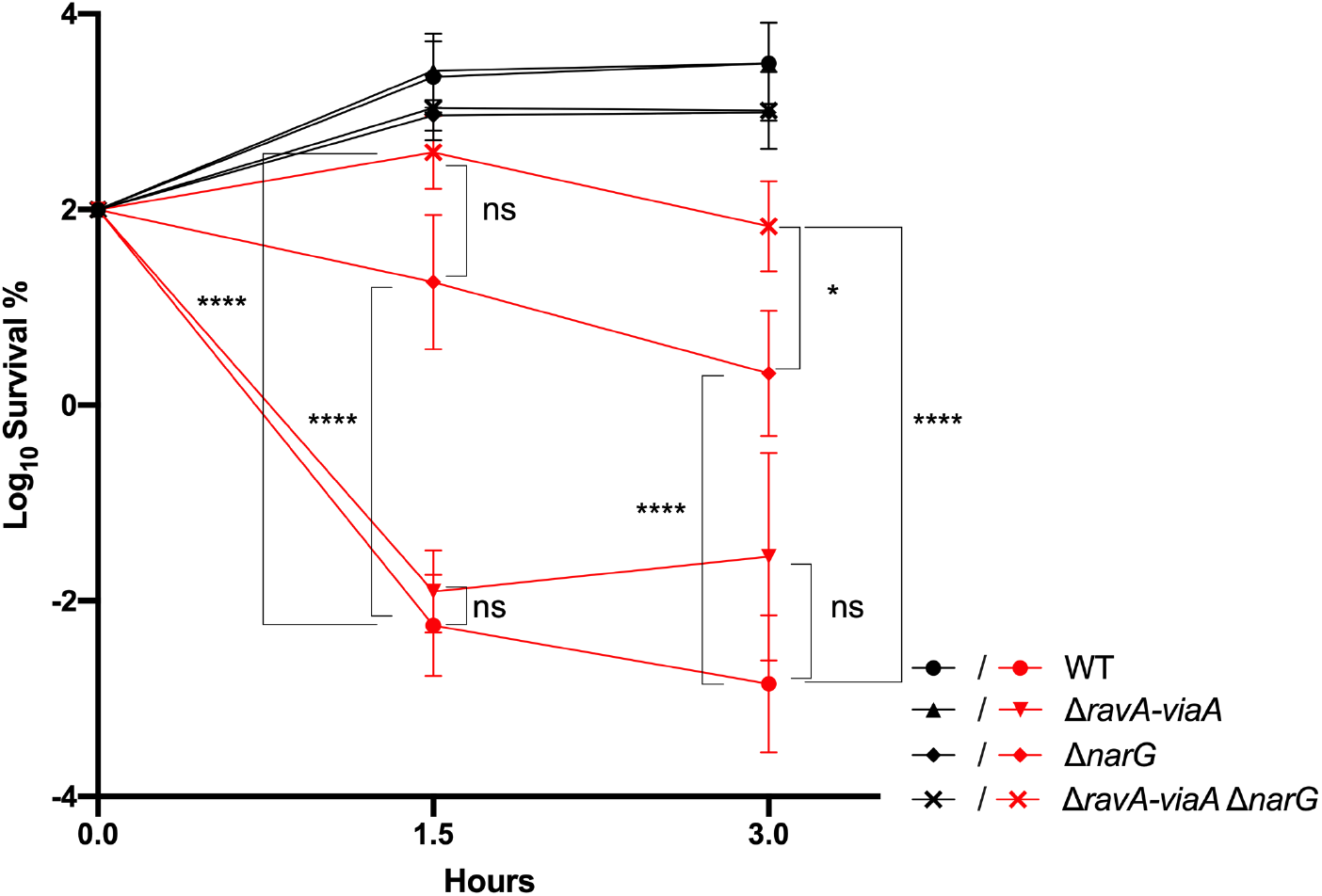
RavA and ViaA do not sensitize nitrate respiring *E. coli* to gentamycin. Survival of WT (FBE051), Δ*ravA-viaA* (FBE706), Δ*narG* (FBE829), Δ*ravA-viaA* Δ*narG* (FBE830) strains after Gm treatment. Cells were grown anaerobically in LB supplemented with nitrate at 10 mM and glycerol at 0.2% and then Gm was added at 16 μg/mL. The survival values after 1.5 and 3 hours of treatment are represented. Black and red lines are for untreated and Gm-treated bacteria, respectively. Lines of untreated WT and Δ*ravA-viaA* (FBE706) strains are overlapping. Lines of untreated Δ*narG* (FBE829) and Δ*ravA-viaA* Δ*narG* (FBE830) are overlapping. Survival measured by CFU per mL, was normalized relative to time zero at which Gm was added (early log phase cells; ~5×10^7^ CFU/mL) and plotted as Log_10_ of % survival. Values are expressed as means of at least 3 biological replicates and error depict standard deviation. One-way ANOVA tests followed by Sidak’s multiple comparaison tests were performed to compare at each time point (1.5 and 3 hours) the treated WT to each of the treated mutant as well as treated Δ*narG* (FBE829) to Δ*ravA-viaA* Δ*narG* (FBE830) (ns: not significant, * adjusted pValue < 0.05 and **** adjusted *p*Value < 0.0001).

### RavA and ViaA sensitize *E. coli* to Gm in the absence of exogenously added electron acceptor in anaerobiosis

Next, we investigated the extant of the RV sensitizing effect in the absence of exogenously added electron acceptor, i.e. in LB medium supplemented, or not, with 0.2% glycerol or 0.2% glucose. First, the MIC values for Gm in all of these media were found to be 8 μg/mL. Next, we performed a killing assay using Gm at 16 μg/mL (2x MIC) in LB (Fig. 3A), in LB glycerol (Fig. 3B) and at 30 μg/mL (~4x MIC) in LB glucose (Fig. 3C). The Δ*ravA-viaA* mutant exhibited increased resistance to Gm compared to the WT strain in all media (Fig. 3). Notably, complementing the Δ*ravA-viaA* mutant with the pRV plasmid suppressed its enhanced sensitivity (Fig. 3 C). These results indicated that RavA ViaA exerts a sensitizing effect on *E. coli* in anaerobic conditions in the absence of exogenous electron acceptor.

**Fig 3.**
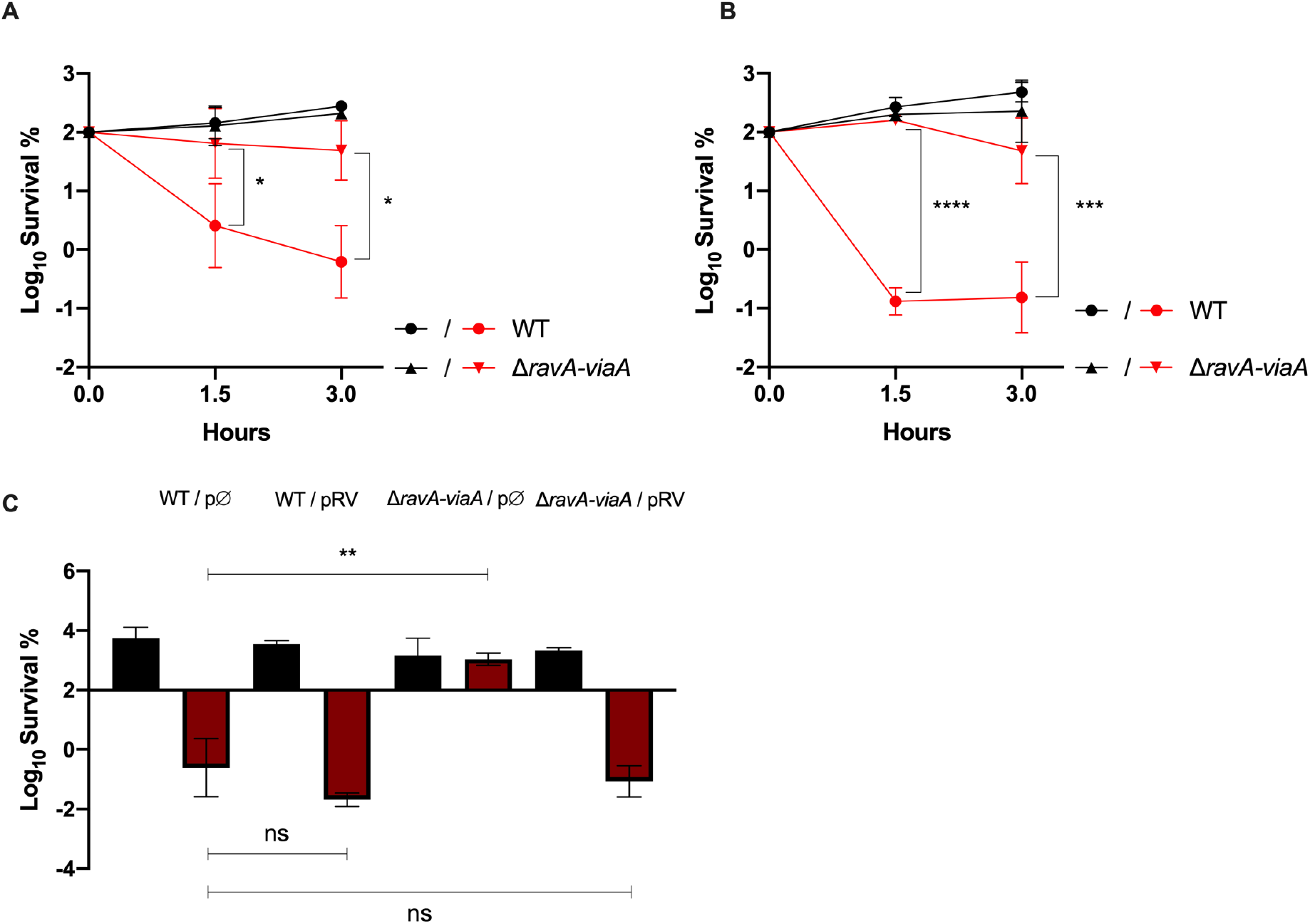
RavA and ViaA sensitize *E. coli* to gentamycin under anaerobic conditions in the absence of exogenous added electron acceptor. (**A and B**) Survival of WT (FBE051) and the Δ*ravA-viaA* (FBE706) strains after Gm treatment. Cells were grown in LB **(A)** or in LB supplemented with 0.2% glycerol **(B)** until OD_600nm_~0.2 and then Gm was added at 16 μg/mL. The survival values after 1.5 and 3 hours of treatment are represented. Black and red lines are for untreated and Gm-treated bacteria, respectively. Values are expressed as means of at least 3 biological replicates and error depict standard deviation. One-way ANOVA tests followed by Sidak’s multiple comparaison tests were performed to compare at each time point (1.5 and 3 hours) the treated WT to Δ*ravA-viaA* mutant (* adjusted pValue < 0.05, *** adjusted *p*Value = 0.0002 and **** adjusted *p*Value < 0.0001). (**C**) Survival of WT (FBE051) and the Δ*ravA-viaA* (FBE706) strains carrying either the pRV plasmid or the control empty vector (pØ). Cells were grown in LB supplemented with glucose (0.2 %), IPTG (1 mM) and ampicillin (50 μg/mL) until OD_600nm_~0.2 and then Gm was added at 30 μg/mL. The survival values after 3 hours of treatment are represented. Black and red bars are for untreated and antibiotic-treated bacteria, respectively. Survival measured by CFU per mL, was normalized relative to time zero at which Gm was added (early log phase cells; ~5×10^7^ CFU/mL) and plotted as Log_10_ of % survival. One-way ANOVA tests followed by Dunnett’s multiple comparaison tests were performed to compare the treated WT to the treated Δ*ravA-viaA* mutant (ns = not significant and ** adjusted pValue < 0.05).

### RavA and ViaA sensitize *E. coli* to aminoglycosides specifically

To test whether the effect of RavA and ViaA proteins is specific to Gm, we measured the survival rate of the Δ*ravA*-*viaA* strain to other antibiotics: another aminoglycoside (Tobramycin), a protein synthesis inhibitor (tetracycline), a fluoroquinolone (Nalidixic Acid) and a β-lactam (ampicillin). We observed that the Δ*ravA-viaA* mutant was more resistant to tobramycin than the WT strain (Fig. 4A). A modest resistance effect was noticed at short time exposure to tetracycline (Fig. 4B). The Δ*ravA*-*viaA* mutation had no effect on the toxic effect of nalidixic acid and ampicillin as its survival rate was the same as the WT strain (Fig. 4C & 4D). Altogether these results showed that in anaerobiosis, RavA and ViaA sensitize *E. coli* specifically to aminoglycosides.

**Fig 4.**
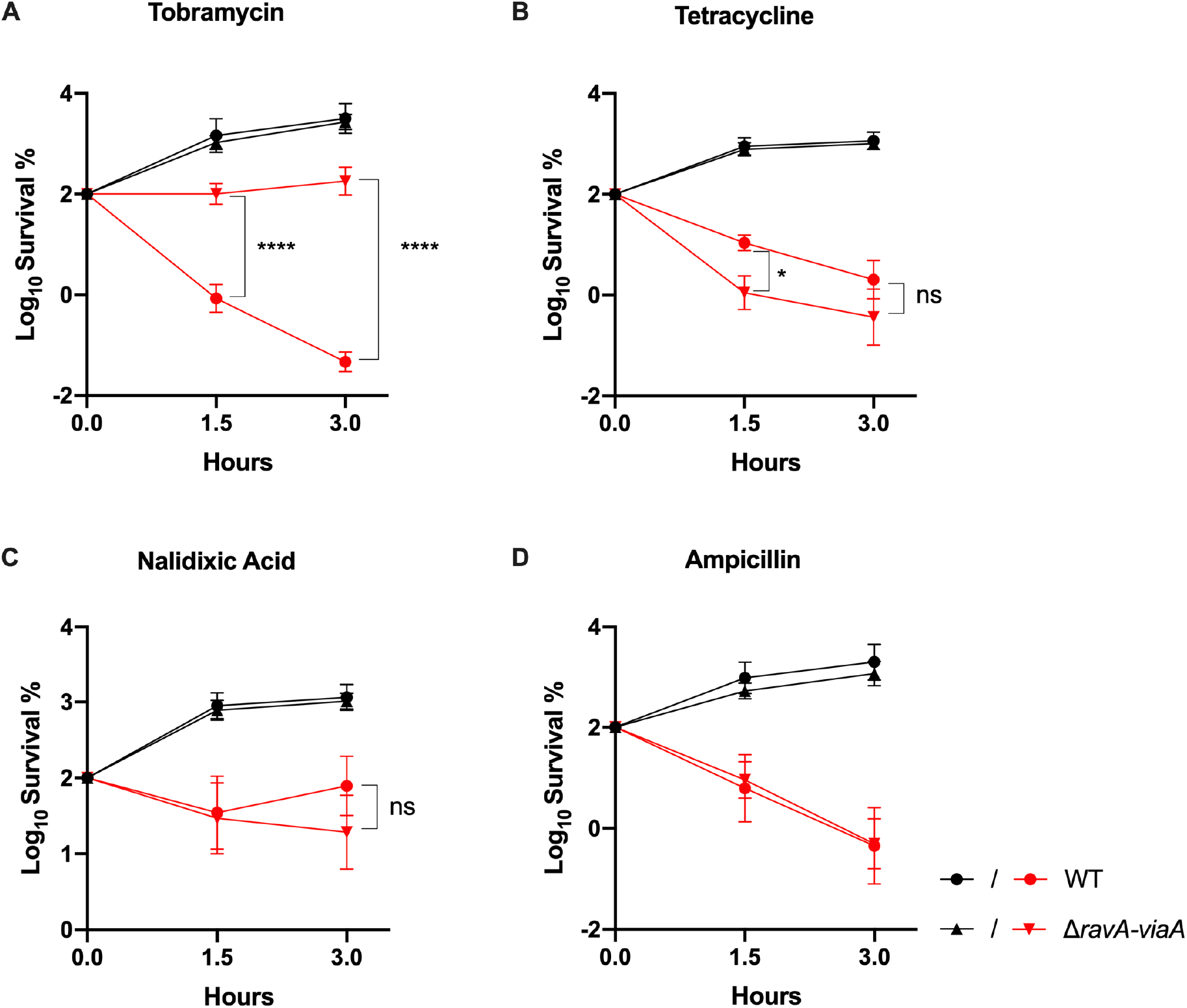
RavA and ViaA sensitize specifically to aminoglycosides. Survival of WT (FBE051) and Δr*avA-viaA* (FBE706) strains after antibiotic treatment. Cells were grown in LB supplemented with glucose (0.2 %) until OD_600nm_~0.1 and antibiotics were added: **(A)** Tobramycin (30 μg/mL); **(B)** Tetracycline (5 μg/mL); **(C)** Nalidixic acid (5 μg/mL); and **(D)** Ampicillin (5 μg/mL). Black and red lines are for untreated and antibiotic-treated bacteria, respectively.The survival values after 1.5 and 3 hours of treatment are represented. Survival, measured by CFU per mL, was normalized relative to time zero at which the antibiotic was added and plotted as Log_10_ of % survival. Values are expressed as means (n=3) and error bars depict standard deviation. One-way ANOVA tests followed by Sidak’s multiple comparaison tests were performed to compare at each time point (1.5 and 3 hours) the treated WT to the treated Δ*ravA-viaA* mutant (ns = not significant, * adjusted pValue < 0.05 and **** adjusted *p*Value < 0.0001).

### RavA and ViaA increase the intracellular gentamycin concentration under anaerobic conditions

To understand the role of RavA ViaA in AG sensitization under anaerobic conditions, we performed a Gm uptake assay using ^3^H-Gm. Gm uptake assay was done with strains that had grown in LB-glucose under anaerobic conditions. Accumulation of ^3^H-Gm in the WT strain progressively increased to reach 1200 ng Gm/10^8^ cells after 2.5 hours (Fig. 5). Instead, in the *ΔravA-viaA* mutant, the accumulation of ^3^H-Gm remained below 100 ng of Gm/10^8^ cells after 2.5 hours. Our results indicated that RavA and ViaA sensitize *E. coli* to Gm by enhancing its uptake and as a consequence its intracellular concentration.

**Fig 5.**
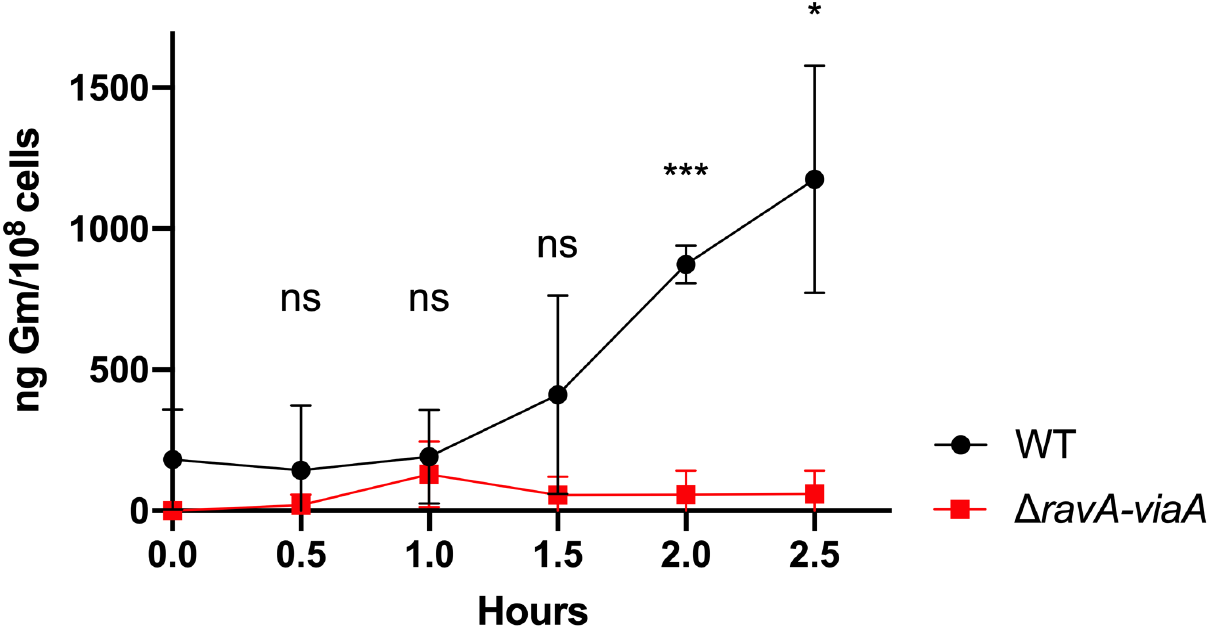
RavA and ViaA increase gentamycin uptake. ^3^H-Gm uptake in WT (FBE051) and Δ*ravA-viaA* (FBE706) strains was measured by incubating early exponential-phase cultures (OD_600nm_~0.1) with 30 μg/mL ^3^H-Gm at 37 °C under anaerobic conditions (LB supplemented with glucose at 0.2%). Values are expressed as means (n=3) and error bars depict standard deviation. Unpaired t-test followed by Welch’s correction was performed to compare the WT strain to the Δ*ravA-viaA* mutant at each time point (ns = not significant, * adjusted pValue < 0.05 and *** adjusted pValue = 0.0003).

### Increased *ravA-viaA* gene dosage enhances aminoglycoside killing of *E. coli* in aerobiosis

The previously reported implication of RV in AG sensitivity was derived from studying *E. coli* grown in the presence of O_2_ [21]. Moreover, this study monitored growth in the presence of sublethal concentrations of kanamycin, another AG. Therefore, we decided to reinvestigate the influence of RV in aerobiosis using killing assays. The MIC of Gm was found to be 2 μg/mL for both wild type (WT) and Δ*ravA-viaA* strains. In a time-dependent killing experiment using a concentration of Gm equivalent to 2.5x MIC (5 μg/mL), *E. coli* WT and Δ*ravA-viaA* strains exhibited similar sensitivities to Gm (Fig. 6A). Hence these results failed to confirm previous data reporting a role of *ravA-viaA* in AG resistance in the presence of O_2_. To further solve this apparent conundrum, we constructed a plasmid carrying the *ravA-viaA* operon (pRV plasmid) and tested whether this would sensitize *E. coli* to Gm. We observed a drastically altered survival of the WT/pRV strain (Fig. 6B) showing the capacity of RavA ViaA to sensitize *E. coli* to Gm killing under aerobiosis. Thus, we concluded that RV can sensitize *E. coli* to Gm in the presence of O_2_ as well but only when they are produced above a threshold level value, that seems not to be reached in exponentially growing cells.

**Fig 6.**
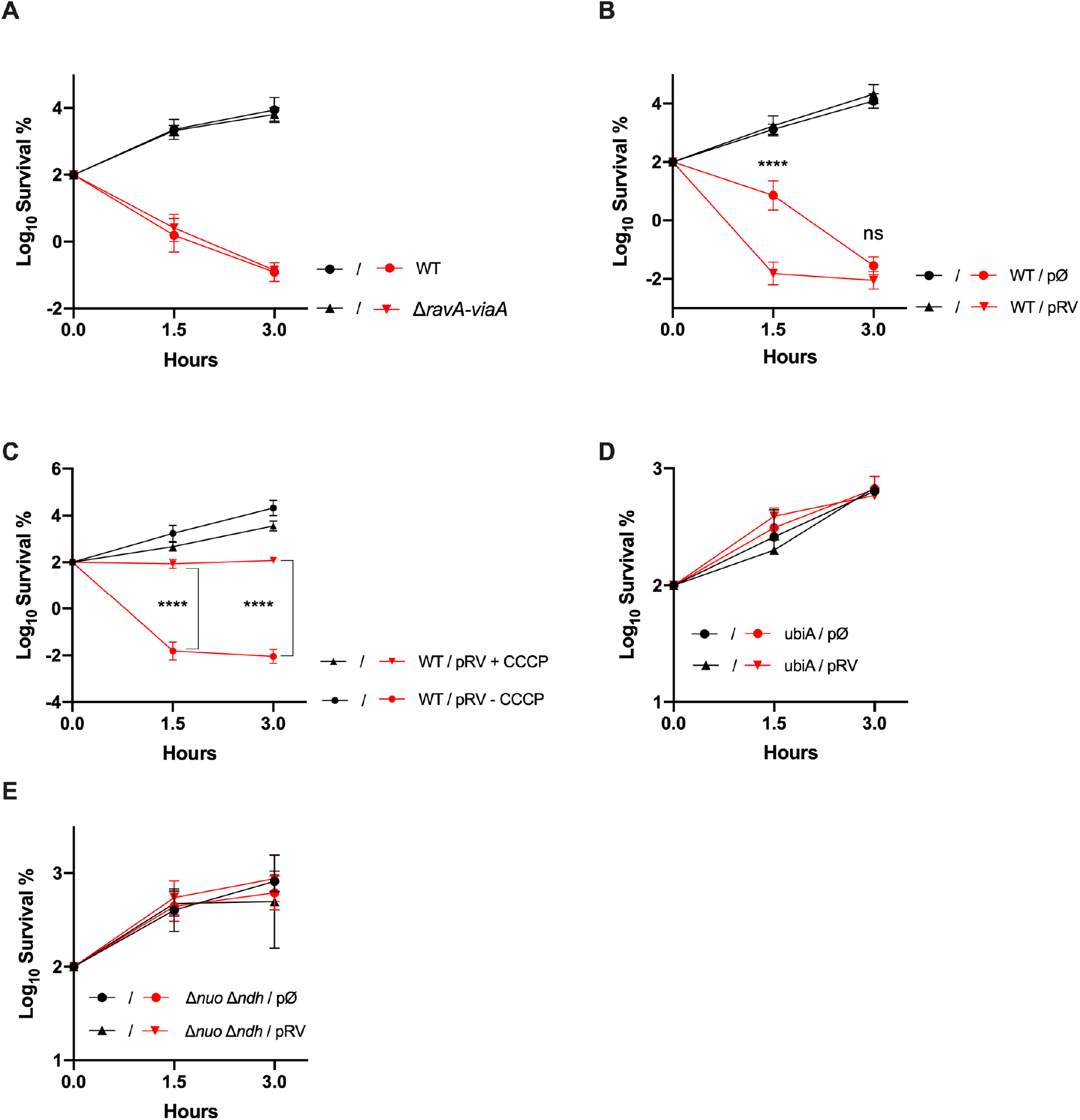
Effect of RavA-ViaA on *E. coli* sensitivity to gentamycin under aerobiosis. **(A-B) Increased *ravA* and *viaA* genes dosage alters survival to gentamycin in killing assay**. Survival of the strain WT (FBE051) and the mutant Δ*ravA-viaA* (FBE706) **(A)** or of the WT (FBE051) strain containing either a plasmid that carries the *ravA-viaA* operon (pRV) or the empty vector control (pØ) **(B)** after treatment with Gm (5 μg/mL) for 1.5 and 3 hours. **(C-E) The RavA-ViaA gentamycin sensitization phenotype is abolished by p.m.f. inhibitor and is dependent upon a functional respiratory chain.** **(C)** Survival of the WT (FBE051) strain containing a plasmid that carries the *ravA-viaA* operon (pRV) after treatment with Gm (5 μg/mL), in the presence or absence of CCCP (5 μg/mL). Survival of Δ*ubiA* (LL922) **(D)** and Δ*nuo* Δ*ndh* (BP1046) **(E)**, containing a plasmid that carries the *ravA-viaA* operon (pRV) or the empty vector control (pØ), after Gm treatment. Cells were grown in LB supplemented with IPTG (1 mM) and ampicillin (50 μg/mL) until OD_600nm_~0.1 and Gm (5 μg/mL) was added. Survival, measured by CFU per mL, was normalized relative to time zero at which the antibiotic was added (early log phase cells; ~5×10^7^ CFU/mL) and plotted as Log of % survival. Values are expressed as means (n=3) and error bars depict standard deviation. Black and red lines are for untreated and Gm-treated, respectively. One-way ANOVA tests followed by Sidak’s multiple comparaison tests were performed (ns = not significant and **** adjusted *p*Value < 0.0001).

### RavA-ViaA mediated *E. coli* killing by aminoglycosides in aerobiosis requires *pmf*

Survival rate to Gm of the WT/pRV was tested in the presence of cyanide-m-chlorophenylhydrazone (CCCP), an ionophore, which dissipates the *pmf*. The data showed that addition of CCCP prevented Gm from killing the WT/pRV strain (Fig. 6C), demonstrating that pRV-mediated killing required *pmf*. Since *pmf* results from respiratory metabolism, we tested whether pRV-mediate sensitization was dependent upon electron transfer chain (ETC)-forming components. We analysed survival to Gm of strains defective for the synthesis of ubiquinones, the lipid that acts as electron carrier within aerobic ETCs. A Δ*ubiA* strain was highly resistant to Gm treatment and pRV plasmid failed to sensitize the strain defective for ubiquinones (Fig. 6D). Similarly, a Δ*nuo* Δ*ndh* strain lacking both NADH dehydrogenase I and II, canceled pRV-mediated sensitization (Fig. 6E). These data supported the notion that respiration is required for pRV-mediated killing in aerobiosis.

### RavA-ViaA have no effect on Nuo activity

To test if RavA and ViaA affect the activity of the Nuo respiratory complex, the Nuo complex activity was measured using deamino-NADH as specific substrate and O_2_ as a final electron acceptor. The level of activity of the Nuo complex was in the Δ*ravA-viaA* mutant comparable whether it carried the pRV plasmid or the empty vector control (S3 Fig). This result led us to conclude that RavA ViaA has no effect on Nuo activity level.

## Discussion

AGs have been used for decades to treat Gram-negative infections. Yet our understanding of their mode of action, and in particular their mode of entry into the cells remains uncertain. A two step model has been put forward, including a first step wherein AG cross the membrane via a *pmf*-dependent pathway followed by a second step wherein aborted translational misfolded polypeptides allow massive entry [27]. In this work, we report the influential role of an AAA+ ATPase from the MoxR subfamily, RavA, associated with its VWA-containing partner, ViaA, in Gm uptake under anaerobiosis. We show that the first *pmf* dependent step is required for RavA-ViaA-dependent facilitating Gm uptake. We discuss a hypothesis wherein RavA-ViaA role would take place when cells are in low energetic state.

*E. coli* possess a highly versatile arsenal of respiratory chains. *E. coli* synthesizes multiple dehydrogenases and terminal reductases, which act as quinone reductases and oxydases, respectively [26,28]. Likewise, from the sole bioenergetic point of view, quinones can connect most of the dehydrogenases with most of the reductases and a great variety of respiratory chains be formed [26,28]. However, not all possible respiratory chain afford redox energy conservation as this requires redox-loop mechanism, which couples electron transfer across the membrane to expulsion of a proton in the periplasm. Energy conservation is maximal in aerobiosis, decreases under fumarate respiration while it gets to its lowest level in the absence of exoneously added electron acceptor [26]. Satisfyingly enough, we found predicted redox energy conservation level to parallel Gm sensitivity level (Fig. 7). Oxygen, nitrate and fumarate as electron acceptors, in this order, yield to higher level of *pmf* and decreasing level of susceptibility of *E. coli* to Gm. Surprisingly, the importance of RavA ViaA in sensitizing *E. coli* to Gm appeared to follow the redox energy hierarchy defined above. Indeed influence of RV was apparent when exogenous electron acceptors providing lowest energy conservation, i.e. no exogenous electron acceptor or with fumarate (Fig. 7). It is unclear which mode *E. coli* is relying on to grow in the absence of “exogeneous added electron acceptor” and we assume that endogenously produced acetate as well as amino acids present in the rich medium can act as electron acceptors. In contrast, with nitrate or oxygen as electron acceptors, which yields to the two highest energy conservation processes, RavA and ViaA had no significant influence. Thus it is tempting to propose as a working hypothesis that RV sensitizing effect takes place only when cell energy goes beyond a threshold value.

**Fig 7.**
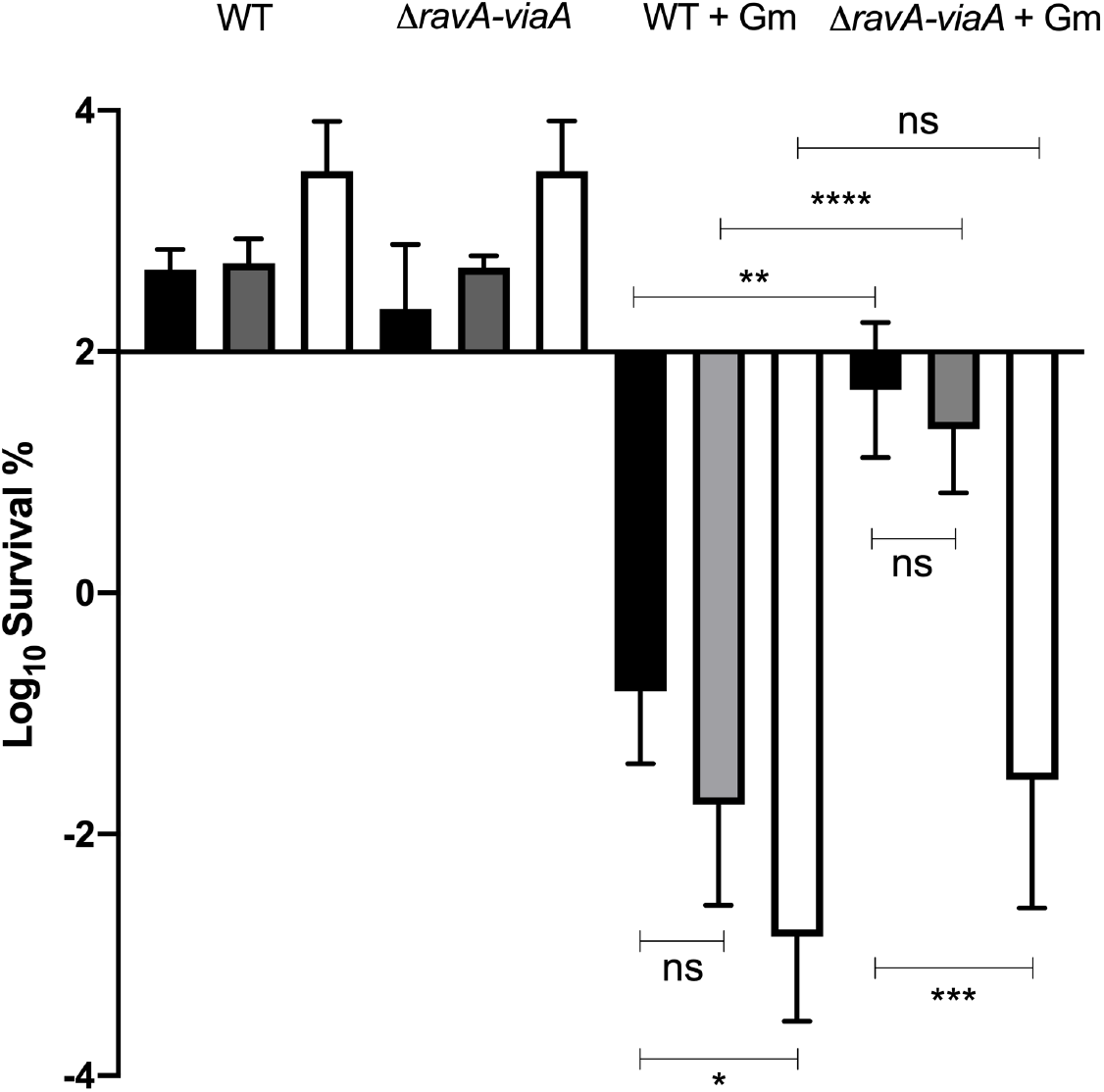
Energy conservation level affects gentamycin sensitivity under anaerobic conditions. Survival of WT (FBE051) and the Δ*ravA-viaA* (FBE706) strains after Gm treatment. Cells were grown in LB-glycerol (black), added with 10 mM fumarate (grey) or with 10 mM nitrate (white) until OD_600nm_~0.2 and then Gm was added at 16 μg/mL. The survival values after 3 hours of treatment are represented. The first two groups represent the untreated strains and the two last groups represent the treated strains (+Gm). Values are expressed as means of at least 3 biological replicates and error depict standard deviation. One-way ANOVA tests followed by Sidak’s multiple comparaison tests were performed (ns = not significant, * adjusted *p*Value < 0.05, ** adjusted *p*Value < 0.005, *** adjusted *p*Value = 0.0003 and **** adjusted *p*Value < 0.0001).

The *ravA-viaA* operon is under Fnr control [19] and consistently we found a phenotype for Δ*ravA-viaA* mutant under anaerobiosis. In contrast, killing assays failed to show evidence of the expected enhanced Gm resistance of *ΔravA-viaA* mutant in the presence of O_2_. Yet our subsequent analysis using plasmid carrying RavA-ViaA copies revealed that RavA ViaA was able to sensitize *E. coli* to Gm, in other words, that aerobic condition was not intrinsically inhibitory to the RavA ViaA activity. Previous study showed the *ravA viaA* operon to be under sigma S control under aerobiosis, with an optimal synthesis of both RavA and ViaA proteins occurring in cells under stationary phase [19]. Hence it is likely that in our killing assays performed on cells growing exponentially in the presence of O_2_, RavA and ViaA protein levels were not high enough for an influence to be detected. It will be interesting to reinvestigate the issue of RavA ViaA influence on Gm lethal activity on stationary resting cells. Besides, if RavA ViaA were to bear a sensitizing effect on starving cells, it would somehow be consistent with the hypothesis above predicting a capacity of RV to sense cells under low energetic metabolism.

We showed that RV complex sensitizes *E. coli* to AG via a *pmf*-dependent mechanism. A simplest explanation to account for the sensitizing role of RV is that RV enhanced respiratory chain activity, thereby yielding to high *pmf* level and enhanced Gm uptake. Yet this explanation seems to fall short. Indeed, no positive effect of RV was found neither on Frd nor on Nuo activity. The effect of RV on Frd was previously investigated and no significant effect was found either [19]. Instead, it was proposed that RV could assist step-wise assembly of the multi-subunits Frd complex in the membrane [19]. Such an hypothesis was in line with the fact that several AAA+ ATPases do assist folding and assembly of multiprotein complexes. Yet, the experiments performed in this study did not show any advantage of the presence of RV on fumarate based growth, casting some doubt on the importance of RavA ViaA for Frd functioning *in vivo*. RavA ViaA were proposed to target the Nuo complex based upon pull-down assays [21]. Actually, only a subset of Nuo subunits did interact with RavA and/or ViaA, and these putative partnerships changed depending upon the growth conditions: NuoA and F interacting with RavA and ViaA under aerobiosis and NuoCD interacting with both RavA and ViaA under anaerobiosis. Yet in their phenotypic analysis, mutating *nuo* genes bore no effect on RavA-ViaA mediated sensitization to sublethal concentration of kanamycin, raising questions about the functional consequences of the interactions between RV and some Nuo subunits by pull-down assays. Moreover, we found no effect of RavA ViaA onto the level of Nuo activity. Thus, if the use of Gm toxicity as a read-out as we used here, clearly demonstrates a physiological link between RV and respiratory complexes, the mode of action of RV on these complexes remains enigmatic.

Antibiotic resistance is a major public health, social and economic problem. AGs are known for their high efficiency against Gram-negative bacteria but their use is restricted to life threatening infections because of their nephrotoxicity and ototoxicity at therapeutic doses [29]. Elucidation of AG sensitization mechanisms in bacteria will allow the use of a decreased effective dose of AGs, to safely treat a wider proportion of infections. Here we identified new molecular actors, RavA and ViaA, which sensitize *E. coli* to AG under anaerobiosis. This is a significant step forward as anaerobiosis is well known to reduce antibacterial activity of AG. This study extands our previous work [4,30,31] and further emphasizes the influence that environmental conditions and composition can bear on level of antibiotic resistance.

## Materials and methods

### Bacterial strains and growth conditions

The *E. coli* K-12 strain MG1655 and its derivatives used in this study are listed in Table 1. Deletion mutations (Δ*ravA*∷Kan^R^, Δ*viaA*∷Kan^R^, Δ*ndh*∷Kan^R^, Δ*frdA*∷Kan^R^, Δ*menA*∷Kan^R^, Δ*nuoC*∷Kan^R^, Δ*glpA*∷Kan^R^, Δ*narG*∷Kan^R^) from the KEIO collection were introduced by P1 transduction. The Δ*ravA-viaA*∷Kan^R^ mutant was constructed using the procedure described by Datsenko K. and Wanner B. [32], using oligos ravAwanner_up and viaAwanner_do, followed by a transduction in clean MG1655 background. Transductants were verified by polymerase chain reaction (PCR), using primers pair hybridizing upstream and downstream the deleted genes. When performed, excision of the kanamycin cassette was done using the pCP20 plasmid [32]. Oligonucleotides used in this study are listed in Table 2. *E. coli* strains were grown at 37°C in Luria-Bertani (LB) rich medium or in minimal M9 medium. Glucose (0.2%), Glycerol (0.2%), IPTG (1 mM), CCCP (5 μg/mL), fumarate (10 mM) or nitrate (10mM) were added when indicated. Solid media contained 1.5% agar. For standard molecular biology techniques antibiotics were used at the following concentrations, kanamycin at 50 μg/mL, and ampicillin at 50 or 100 μg/mL.

**Table 1.** *E. coli* strains used in this study.

| *E. coli* strains | Relevant genotype | Source |
| --- | --- | --- |
| BP897 | MG1655 $\Delta nuo::nptI$ Kan<sup>R</sup> | Ezraty B. *et al.* [4] |
| BP1046 | MG1655 $\Delta nuo::nptI$ $\Delta ndh::FRT$ | This study |
| FBE051 | MG1655 Wild type | Lab collection |
| FBE501 | MG1655 $\Delta menA::FRT$ | This study |
| FBE706 | MG1655 $\Delta ravA-viaA::FRT$ | This study |
| FBE790 | MG1655 $\Delta frdA::FRT$ | This study |
| FBE829 | MG1655 $\Delta narG::FRT$ | This study |
| FBE830 | MG1655 $\Delta ravA-viaA$ $\Delta narG::FRT$ | This study |
| FBE831 | MG1655 $\Delta ravA-viaA$ $\Delta frdA::FRT$ | This study |
| FBE950 | MG1655 $\Delta glpA::FRT$ | This study |
| FBE951 | MG1655 $\Delta ravA-viaA$ $\Delta glpA::FRT$ | This study |
| FBE1055 | MG1655 $\Delta glpA$ $\Delta nuoC::FRT$ | This study |
| FBE1057 | MG1655 $\Delta nuoC::FRT$ | This study |
| LL922 | $\Delta ubiA$ | Kazemzadeh K. *et al.* [34] |

**Table 2.**
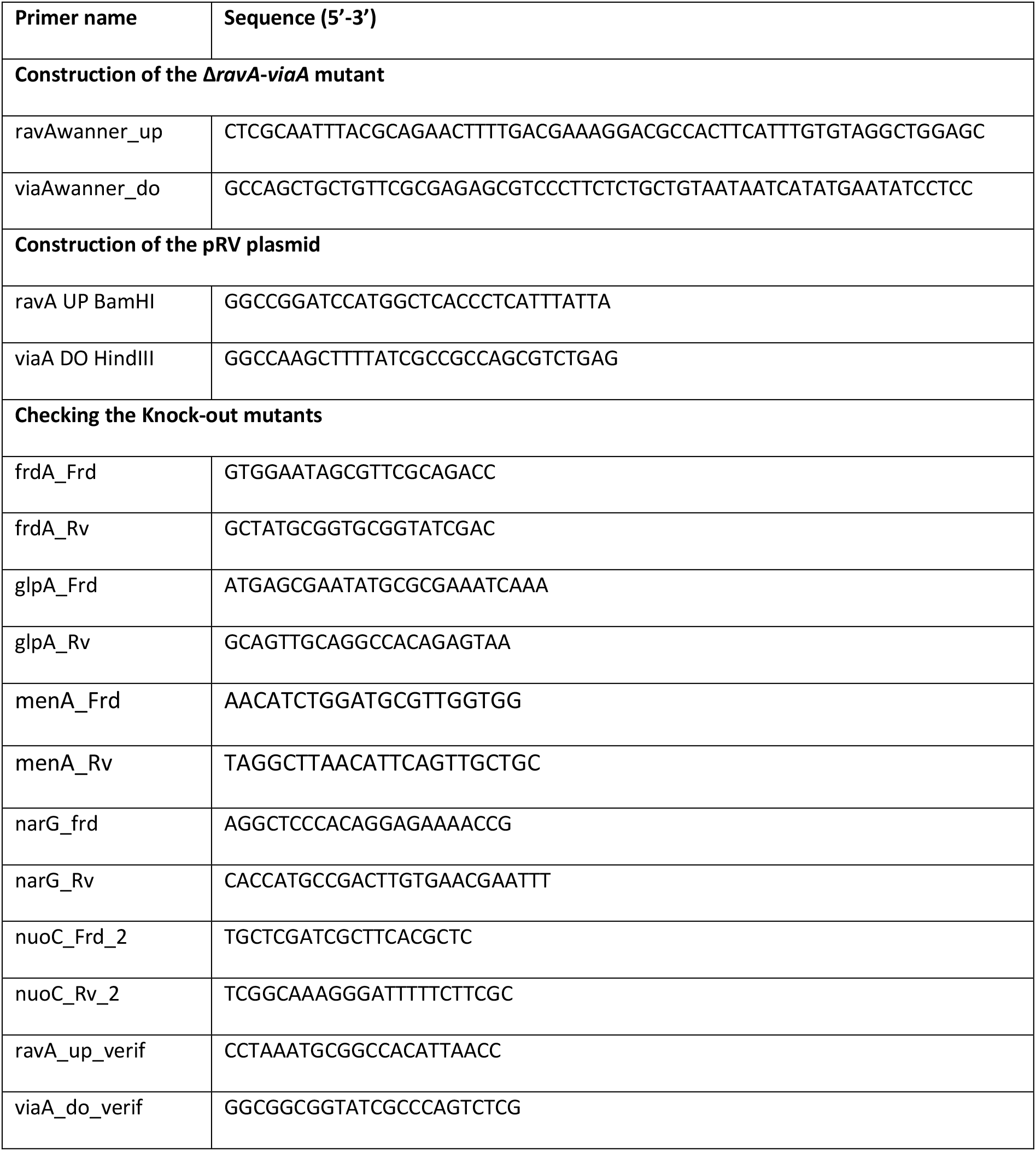
Oligonucleotides used in this study.

### Plasmid construction

Plasmid pRV was constructed by, first, PCR amplification of the coding region of *ravA-viaA* from the *E. coli* MG1655 chromosomal DNA using the following primers pair: ravA UP BamHI/ viaA DO HindIII (Table 2). The PCR product was then digested by *BamH*I and *Hind*III and cloned into the *BamH*I/*Hind*III linearized ptrC99A vector [33]. The sequence of the inserted fragment was checked by DNA sequencing.

### Time-dependent killing assay

Overnight cultures were diluted (1/100) and grown aerobically or anaerobically in specific medium as indicated in the figures’ legends at 37°C to an OD_600_ of 0.2. At this point (T0) antibiotics at the indicated concentration were added to the cells. At different incubation times, 100 μL of cells were diluted in sterile phosphate buffered saline solution (PBS buffer), spotted on LB agar and then incubated at 37°C for 24 to 48h. Cell survival was determined by counting colony-forming units per mL (CFU/mL). The absolute CFU at time-point 0 (used as the 100%) was ≈ 5×10^7^ CFU/mL. Survival rate in anaerobic conditions was performed in anaerobic chamber (Coy and Jacomex Chambers). Materials (medium, tubes, plates…) were all previously equilibrated in the anaerobic chamber for at least 18 h.

### MIC determination

The MICs were determined by the microdilution method in a 96 wells plate according to the Clinical Laboratory Standards Institute (CLSI) guideline. Briefly, serial dilutions of Gm in a 2-fold manner were done in 100 μL Cation Adjusted Müller-Hinton or in LB supplemented or not with either glucose (0.2%) or fumarate (10 mM) or nitrate (10mM). *E. coli* inoculum were prepared by suspending colonies grown overnight on LB agar using 1×PBS to achieve a turbidity of 0.5 McFarland (1 × 10^8^ CFU/ml) and the final concentration of the inoculum in each well was around 5 × 10^5^ CFU/mL. The plates were incubated at 37°C for 18 hours under aerobic or anaerobic conditions. MIC was defined as the lowest drug concentration that exhibited complete inhibition of microbial growth. All MICs were determined from at least three independent biological replicates.

### Gentamycin uptake

[^3^H]-Gm (20 μCi/mg; Hartmann Analytic Corp.) was added at the indicated final concentration and cultures were incubated at 37°C on a rotary shaker. At given times, 500 μL aliquots were removed and collected on a 0.45 μM-pore-size HAWP membrane filter (Millipore) pre-treated with 1 mL of unlabelled Gm (250 μg/mL). Filters were subsequently washed with 10 ml of 3 % NaCl, placed into counting vials, dried for 30 min at 52°C whereafter 8 mL of scintillation liquid were added and incubated overnight at room temperature. Vials were counted for 5 min. Gm uptake efficiency is expressed as total accumulation of Gm (ng) per 10^8^ cells.

### Competition experiment in batch culture

The two strains tested were first grown separately overnight in M9 medium supplemented with casamino acids (0.1%). The cell density of each suspension was measured by OD_600nm_ reading and by CFU count. Each overnight culture containing approximately 3×10^8^ cells/mL was diluted 1/100-fold and mixed in a ratio of 1:1 to inoculate 25 mL of M9 supplemented with casamino acids (0.1%) (time 0 h) and incubated for 24 h at 37°C for a competitive growth. The co-culture was diluted 1/100 in 25 ml of fresh M9 medium and grown for another 24 h at 37°C. The initial density of each strain was determined in the initial co-culture (0 h) from CFU data by diluting and plating population samples onto LB agar and LB agar supplemented with kanamycin. Similarly, the final density of each strain was determined for all the co-cultures.

### Enzymatic assay

Cells grown in LB (100 mL) to OD_600nm_ 0.6 were harvested by centrifugation, whashed once in 50 mM phosphate buffer pH 7.5, and resuspended in 50 mM phosphate buffer pH 7.5 (6 mL), lysed using a French press, aliquoted (100 mL) and frozen immediately in liquid nitrogen. Nuo activity was assayed at 30°C by adding thawed samples to 50 mM phosphate buffer pH 7.5 containing reduced nicotinamide hypoxanthine dinucleotide (deamino-NADH) (250 mM) as specific substrate, and by following A_340nm_. Protein concentration was determined using the protein A_280nm_ method on NanoDrop2000 spectrophotometer.

#### Acknowledgments

We thank all members of the Py group (Marseille), the Barras unit (Paris) and Irina Gutsche (IBS, Grenoble) for fruitful discussions.

#### Authors Contributions

**Conceptualization**: Frédéric Barras, Béatrice Py.

**Funding acquisition**: Frédéric Barras.

**Investigation and methodology**: Jessica Y. El Khoury, Jordi Zamarreño Beas, Allison Huguenot, Béatrice Py.

**Supervision**: Frédéric Barras, Béatrice Py.

**Writing review & editing**: Frédéric Barras, Béatrice Py, Jessica Y. El Khoury.

## Supporting information

**S1. Fig.**
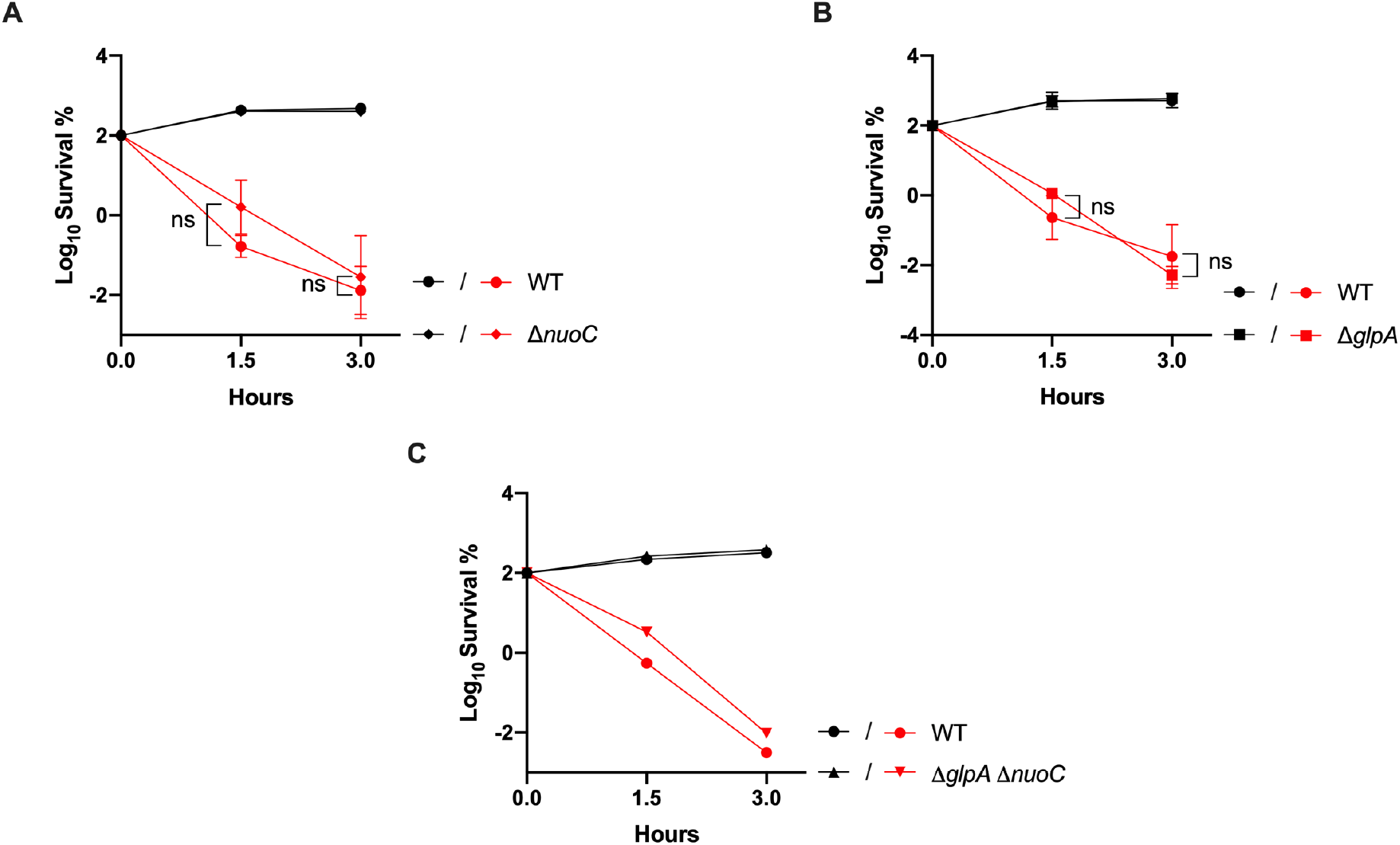
The Nuo complex and the GlpA complex are dispensable for the RavA/ViaA-dependent sensitization of *E. coli* to gentamycin under fumarate respiration. **(A, B, C)** Survival of WT (FBE051), Δ*nuoC* (FBE1057), Δ*glpA* (FBE950) and Δ*glpA* Δ*nuoC* (FBE1055) strains after Gm treatment. Cells were grown in LB supplemented with fumarate at 10 mM **(A)** and glycerol at 0.2% **(B, C)** and then Gm was added at 16 μg/mL. The survival values after 1.5 and 3 hours of treatment are represented. Black and red lines are for untreated and Gm-treated bacteria, respectively. Most of the lines of untreated cells are overlapping. Survival measured by CFU per mL, was normalized relative to time zero at which Gm was added (early log phase cells; ~5×10^7^ CFU/mL) and plotted as Log of % survival. For **A** and **B**, values are expressed as means of at least 3 biological replicates and error depict standard deviation. One-way ANOVA tests followed by Sidak’s multiple comparaison tests were performed to compare at each time point (1.5 and 3 hours) the treated WT to each of the treated mutant (ns = not significant).

**S2 Fig.**
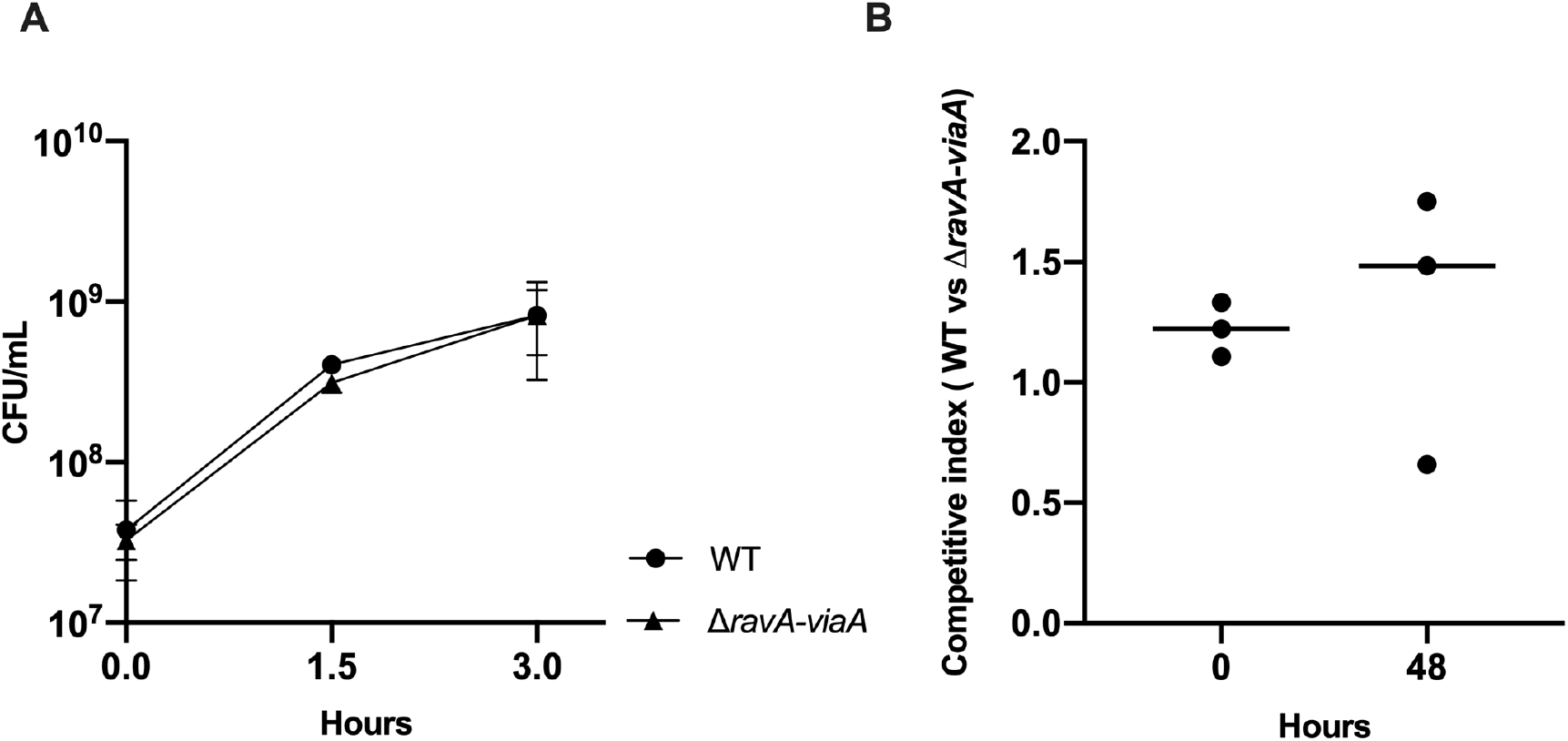
RavA-ViaA have no effect on fumarate respiration dependent growth. (**A**) CFU/mL of WT (FBE051) and Δ*ravA-viaA* (FBE706) strains were determined when grown anaerobically in LB medium supplemented with fumarate (10 mM). At time 0, the OD_600nm_ of the culture was approximately 0.1. (**B**) Strains WT (FBE051) and Δ*ravA*-*viaA* (FBE706) were grown separately overnight in minimum M9 medium supplemented with glycerol (0.2 %), fumarate (10 mM) and casamino acids (0.1 %), in anaerobic conditions. Cultures were then diluted 1/100 into fresh medium and co-inoculated in a 1:1 ratio (t_0_). The co-culture was incubated for 48h at 37°C for a competitive growth. The competitive index was calculated as follow (CFU_mutant_/CFU_wt_)t_48_/(CFU_mutant_/CFU_wt_)t_0_ (circles represent the values obtained in three independent experiments, n=3, lines represent the medians).

**S3 Fig.**
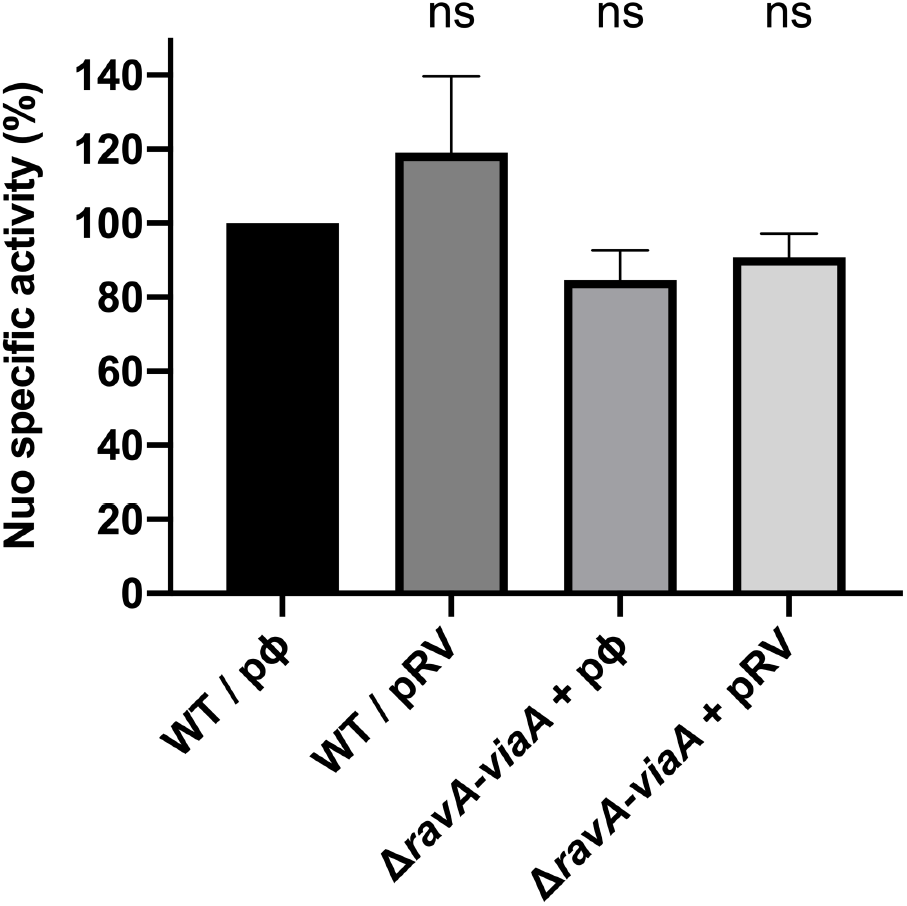
RavA-ViaA have no effect on Nuo activity. Nuo specific activity in the WT (FBE051) and the Δ*ravA-viaA* (FBE706) strains containing the plasmid carrying the *ravA-viaA* genes (pRV) or the corresponding empty vector (pØ). Nuo specific activity was measured in cells extracts using deamino-NADH as substrate. Values are expressed as means (n≥3) and error bars depict mean deviation. One-way ANOVA tests followed by Dunnett’s multiple comparaison tests were performed (ns = not significant). The 100 % correspond to the activity in the WT strain is 127 nmol/min/mg protein.

